# Non-local model of chemotaxis based on peer attraction

**DOI:** 10.1101/2023.05.05.539547

**Authors:** Lionel Dupuy, Matthias Mimault, Mariya Ptashnyk

## Abstract

Movement is critical for bacterial species inhabiting soils because nutrient availability is limited and heterogeneously distributed both in space and time. Recent live microscopy experiments show that bacteria form flocks when navigating through porous medium, and complex cell-cell interactions may be required to maintain such flocks. Here we propose a non-local model to study how peer attraction can affect flocking patterns in a porous medium. We establish the existence and uniqueness of the solution of the problem, propose a numerical scheme for simulations of the non-local convection-diffusion equation, and investigate the numerical convergence of the scheme. Numerical simulations showed that the strength of peer attraction is critical to control the size, shape, and nature of movement of the flocks in a porous network.

**MSC Classification:** 35F31, 92Cxx, 92-10

## 1 Introduction

Soil bio-available carbon is limited and distributed heterogeneously both in space and time [1]. Soil microorganisms, occupying an insignificant fraction of the available space [2], must be able to sense the presence of nutrients in their environment to move towards patches of nutrient and proliferate. The mobility of individual bacterial cells can be observed in laboratory conditions, and the mechanisms involved in movement in liquids or on solid surfaces have been well described. These include flagella mediated swimming, twitching, swarming, or passive dispersion [3]. Bacteria also possess numerous receptors which enable them to detect distant nutrient sources and direct their movement to colonise regions of soils where nutrients are present, e.g. plant roots [4]. The detailed knowledge of bacterial motility in laboratory conditions has logically resulted in the formulation of models for bacterial dynamics, e.g. the Keller-Segel model for chemotaxis [5], which can reliably predict bacterial proliferation patterns observed in laboratory conditions [6].

The movement of bacteria in soil, however, is poorly characterised. Natural soils are opaque. Non-destructive live imaging techniques cannot easily resolve microbial structure and data acquisition time is too slow to resolve bacterial movements [8]. The soil structure is complex and it affects the mobility of bacteria. Not all the pores are accessible to biological cells because of size incompatibility, and the connectivity of accessible pores determine the paths through which a bacterial cell can move towards a nutrient source [9]. Soil water content also affects the connectivity of soil pores. When saturated, bacterial cells can move through the pore freely, but at lower soil water content, movement is limited to water films at the surface of particles [10, 11]. It has been shown that soil water content relates directly to the speed of travel of various bacteria through soil [12]. The dynamics and distribution of soil nutrients are also complex and heterogeneous [13], and it is not clear which chemical signals are perceived and exploited by soil bacteria.

Usually, mathematical models for bacterial movement require coupled systems of partial differential equations to describe on one hand the evolution of the nutrient sources acting as signals, and on the other hand the growth and chemotactic movement of the population of bacteria towards the chemical signals [14, 15]. Such chemotaxis models have been calibrated on soil data, but large variations in estimated chemotaxis parameters indicates that the models may be inadequate [16]. Recent progress in live microscopy within granular materials [17] have also confirmed that bacteria migrate as flocks to move efficiently through the soil pore structure [7], and coordination through complex cell-cell interactions may be needed, see Fig. 1. There has been extensive work to improve models for bacterial movement, for example coupling the dynamics of the fluid to swimming of bacteria [18], but these models require many parameters and it is difficult to incorporate the soil microstructure.

**Fig. 1:**
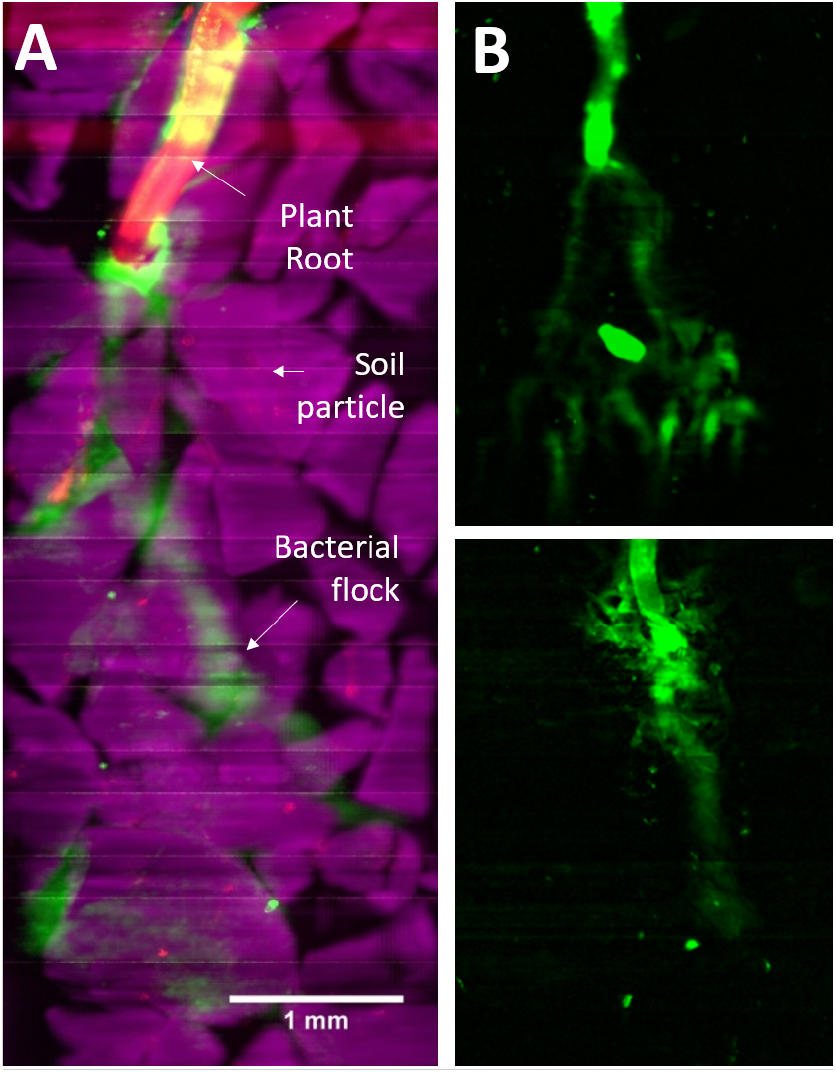
Bacterial flocks observed in soil [7]. (A) Bacterial flocks (green) are observed forming between soil particles (magenta) and moving towards the tip of a plant root (red). (B) The morphology and dynamics of the flocks is affected by biological factors but also by the external environment. When the soil solution has higher viscosity (top), the flocks are more branched and have thinner diameter than when the soil solution has the viscosity of water (bottom).

To investigate how peer attraction may regulate the formation of bacterial flocks observed during the colonisation of plant roots, we propose to develop non-local approaches to model bacterial movement in soil. Non-local models have successfully described complex interactions inside populations of agents such as crowds of pedestrian, schools of fish, flocks of bird, and colonies of bacteria [19–21]. Here, we formulate a model that determines the velocity of bacterial cells as a function of the strength of cell-cell interactions, dispersion and the chemotactic potential determined by the presence of the plant root in the soil domain. We establish the existence and uniqueness results for the new model and convergence properties of the proposed numerical scheme. Numerical simulations show that the non-local model reproduces the flocking patterns observed experimentally, with a reduced number of parameters, and identify factors affecting the morphology and dynamics of the flocks.

## 2 Derivation and analysis of the mathematical model

We consider a two dimensional Lipschitz domain Ω ⊂ ℝ^2^, representing the soil pore space around a plant root. The transport of bacterial cell density *b*, defined as the number of bacterial cells per unit volume, between soil particles is considered to depend on the attraction by the plant root, and peer attraction between bacteria. Then the model for the bacterial cell density dynamics reads

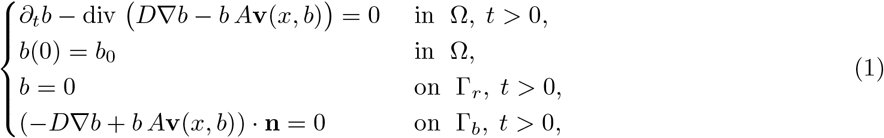

where *A* is the speed of the bacterial cells, **v** determines the direction of the velocity of bacterial cells, and *D >* 0 is the diffusion coefficient. Here Γ_*r*_ ⊂ *∂*Ω denote the part of the plant root surface colonised by bacteria and Γ_*b*_ = *∂*Ω\ Γ_*r*_. The model incorporates the microstructure of soil where particles are represented as obstacles to the movement of bacterial cells. The Neumann boundary conditions on Γ_*b*_ represent the fact that bacteria are blocked by the soil particles and external boundaries of domain Ω. The zero Dirichlet boundary conditions on Γ_*r*_ represent the colonisation of the root and bio-film formation. The vector field **v** is given by

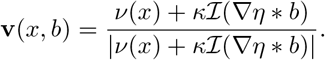

It is a function of the gradient of a chemo-attractant exuded by the root *ν* : Ω → ℝ^2^ and the peer attraction between bacteria of strength *κ* and direction ℐ. We model the movements of bacteria towards the vicinity of the root which chemical distribution is assumed at steady state. This movement is assumed to be fast so that bacterial populations do not have the time to affect the chemical gradients in soil. Hence, the distribution of the chemo-attractant concentration is assumed to be independent of time. We define *ν* as the geodesic field oriented towards the root which can be determined as the gradient of the distance map from the root *φ*, i.e. *ν* = ∇*φ*, where *φ* is the solution of the eikonal equation

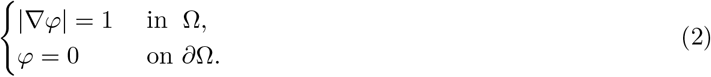

The direction of the movement due to peer attraction ℐ is determined based on the perception of the bacterial cell density in a neighbourhood. This is represented by a smooth non-negative convolution kernel *η* : ℝ^2^ → ℝ with compact support

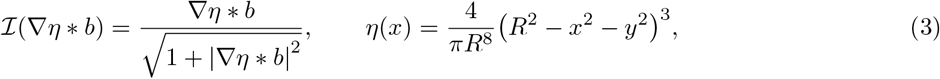

where |·| denotes the Euclidean norm and *R >* 0 is the interaction radius. To define the convolution ∇*η* ∗ *b* for all *x* ∈ Ω, we extend *b* so that *b*(*t, x*) = 0 for 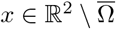 and *t >* 0.

For initial condition we consider *b*_0_ ∈ *L*^2^(Ω).

### Definition 1.

*A weak solution of problem* (1) *is a function* 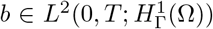, *with* 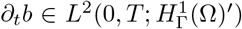, *satisfying*

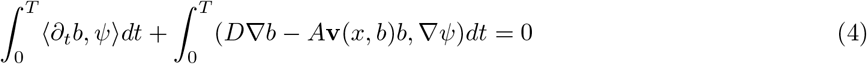

*for any* 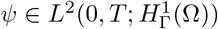, *and initial condition b*(0) = *b*_0_ *in the L*^2^*-sense*.

Here ⟨·, ·⟩ and (·, ·) denote the dual product between 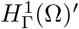 and 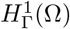 and the scalar product in *L*^2^(Ω), respectively, and

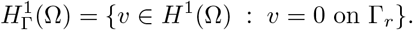

The approach to prove the well-posedness of model (1) is to linearise the equations before applying the Banach fixed-point theorem, which then yields the existence of a unique solution. For this, we first derive a priori estimates for solutions of (1).

### Lemma 1.

*Solutions of* (1) *satisfy the following a priori estimates*

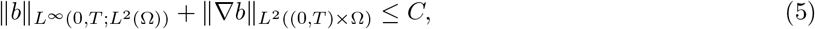

*for some constant C >* 0, *depending on A, D, initial condition b*_0_, *and T*. *Proof*. Considering *b* as a test function in (4) yields

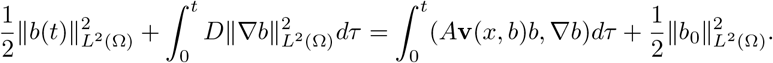

Since ∥**v**∥_*L∞*((0,*T*)×Ω)_ ≤ 1, applying the Hölder inequality yields

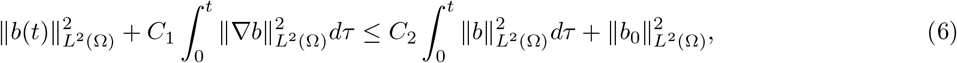

with *C*_1_ = 2*D* − *δA* and *C*_2_ = *A/δ* for any 0 *< δ < D/A*. Then Gronwall’s inequality implies

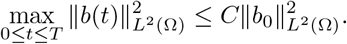

Using the last estimate in (6) we then obtain

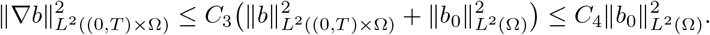

Combining the last two estimates yields a priori estimates stated in the lemma.

### Theorem 1.

*For b*_0_ ∈ *L*^2^(Ω), *there exists a unique weak solution of problem* (1). *Proof*. For a given 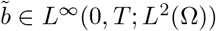 ∈ *L*^∞^(0, *T* ; *L*^2^(Ω)), we first consider the linear problem

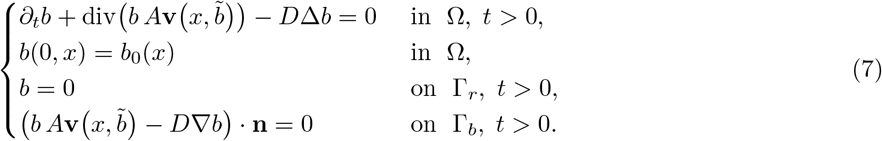

Notice that there exists a unique viscosity solution 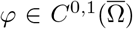 of (2), see e.g. [22, 23]. Hence, together with 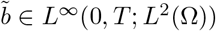, the Vector field 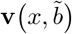 is well-defined in *L*^∞^(0, *T* ; *L*^∞^(Ω)^2^).

The existence of a solution of (7) can be shown using the Galerkin method and a priori estimates similar to those in Lemma 1, derived for the Galerkin approximation.

Next, we apply the Banach fixed-point theorem to show existence of a unique solution of the nonlinear problem (1). To show the contraction properties for the map 𝒦 : *L*^∞^(0, *T* ; *L*^2^(Ω)) → *L*^∞^(0, *T* ; *L*^2^(Ω)), where for 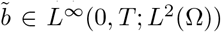 we define 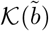 as solution of (7), we consider problem (7) for given 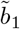 and 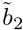, together with their associated solutions *b*_1_ and *b*_2_. Then for the difference *b*_1_ − *b*_2_ we obtain

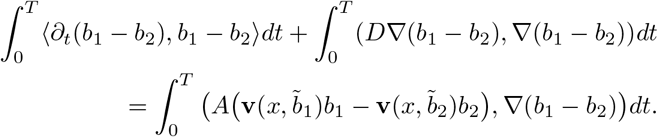

First we show that **v**(*x, b*) is Lipschitz-continuous with respect to *b*. Consider

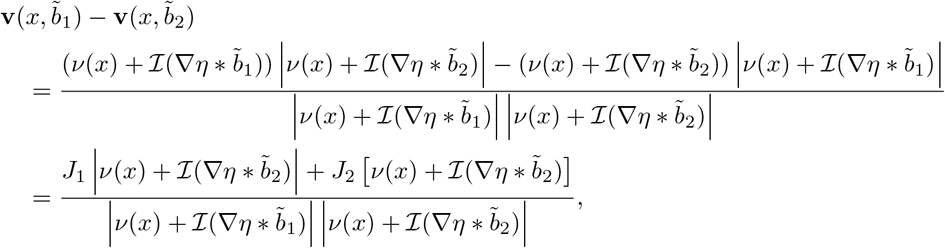

where 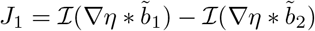 and 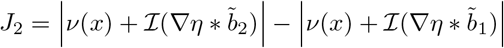. From the definition of ℐ(∇*η* ∗ *b*) we have

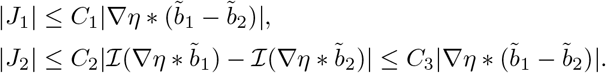

Hence, we obtain

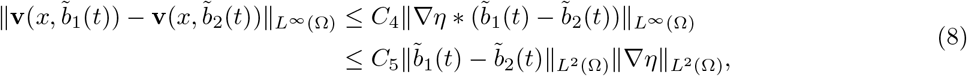

for *t* ∈ (0, *T*]. Using (8) and the boundedness of **v** yields

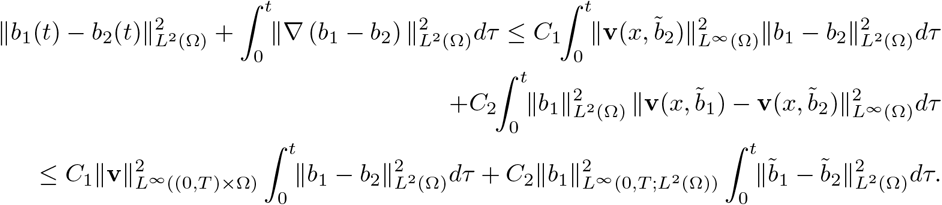

By application of Gronwall’s inequality, we obtain

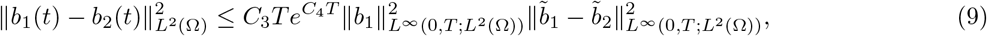

for *t* (0, *T*]. Thus for sufficiently small time interval *T*, such that 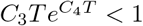, the operator 𝒦 is a contraction. Applying the Banach fixed-point theorem implies the existence of a unique solution of the nonlinear problem (1). Since the time interval depends only on the model parameter and is independent of the solution, we can iterate over the time interval and obtain the existence of a unique solution for any *T >* 0.

## 3 Numerical scheme for the model

For the numerical simulation of model (1), we consider a Cartesian discretisation of domain Ω⊂ ℝ^2^ and the central finite-difference method with dimensional splitting [24], with alternative updates along each space variable *x*_1_ and *x*_2_

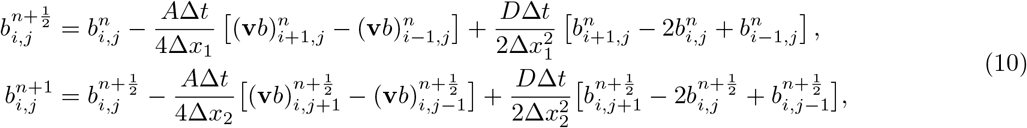

where 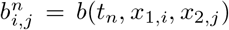 and *x*_1,*i*_ = *i*Δ*x*_1_, *x*_2,*j*_ = *j*Δ*x*_2_, for *i* = 1, …, 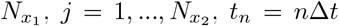, for *n* = 1, …, *N*_*t*_. We consider 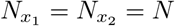 and Δ*x*_1_ = Δ*x*_2_ = Δ*x* satisfying

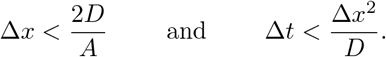

Additionally, the time step is linked to the space step through the Courant-Friedrichs-Levy constant *λ*_CFL_ such that Δ*t* = *λ*_CFL_Δ*x*^2^, with *λ*_CFL_ ≤ 1*/D*.

The discretisation of the velocity vector is given by

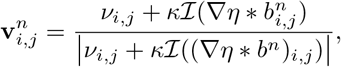

where the convolution is computed through the following quadrature expression

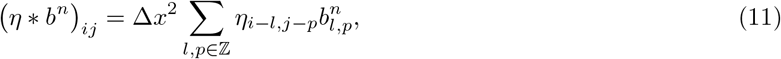

assuming 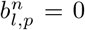 for the grid points outside Ω, and implemented using the fast 2-D convolution function in Matlab [25]. The discrete vector field *ν*_*i,j*_ is generated by solving numerically the eikonal equation (2) using the fast sweeping method [26]. It approximates the solution in an iterative way by sweeping the domain several times along each space direction and updating the solution values to satisfy

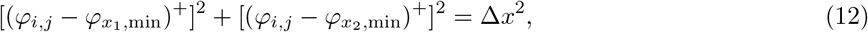

where 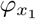,_min_ = min(*φ*_*i*−1,*j*_, *φ*_*i*+1,*j*_), 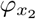,_min_ = min(*φ*_*i,j*−1_, *φ*_*i,j*+1_), (*v*)^+^ = max{*v*, 0}, and *i, j* vary between 2, *N* − 1 in the following four different orders, depending on the direction of the sweeping of the domain Ω,

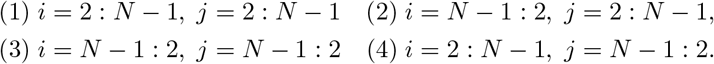

The boundary conditions are fixed to *φ* = 0 at the objective boundary, from where the distance is computed, and a sufficiently large value anywhere else on the boundary. Here, the objective boundary is Γ_*r*_, the surface of the plant, and the boundary condition is set to *φ* = *φ*_max_, which is twice the maximal distance, on Γ_*b*_. The approximated solution of (2) is initialised by *φ*^0^ = *φ*_max_ on Ω*/∂*Ω. The update of the approximation is then given by

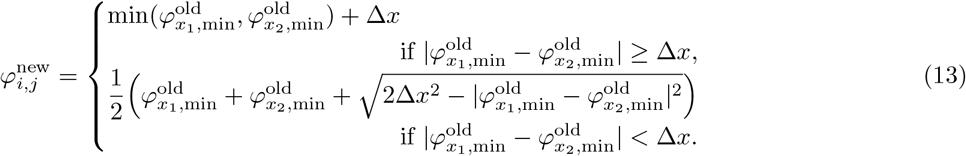

The iteration and update of the approximation of *φ* stops when the *L*_2_-norm of 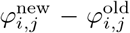 falls below a tolerance criteria *ε* = 10^−3^.Then the discrete velocity *ν*_*i,j*_ is obtained from the approximation of the gradient of *φ*, accounting for the presence of obstacle (here soil particles) in Ω. For each grid point inside the domain Ω, we use the central difference, whereas for the points adjacent to *∂*Ω, we use the forward and backward difference depending on the direction of the normal vector to *∂*Ω. Thus *ν*_*i,j*_ = (∇*φ*)_*ij*_, with

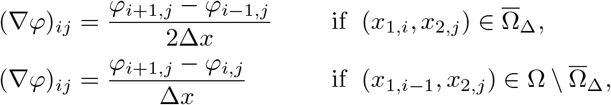

where Ω_Δ_ = {*x* ∈ Ω : dist(*x, ∂*Ω) *>* Δ*x*}, see Figure 2 for illustration.

**Fig. 2:**
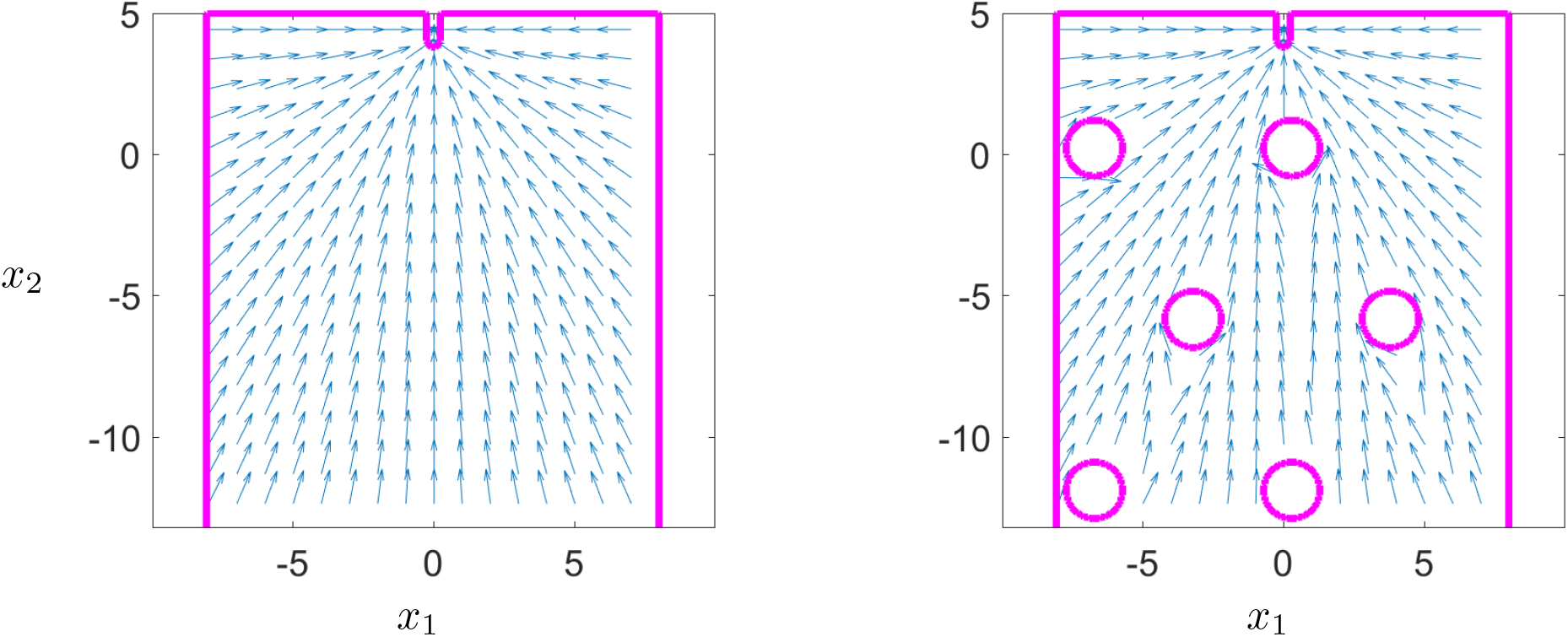
Illustrations of the vector fields ∇*φ* without (left) and with (right) obstacles on the domain Ω = (−8.0, 8.0) × (−13, 5).

### Order of convergence

To estimate the order of convergence of the numerical scheme (10), (12), we consider the analytic solution *b*(*t, x*_1_, *x*_2_) = cos(*x*_1_) sin(*x*_2_)*e*^*t*^ for (*x*_1_, *x*_2_) Ω = (2*π*, 2*π*) (2*π*, 2*π*) and *t >* 0, of the equivalent problem with a source term

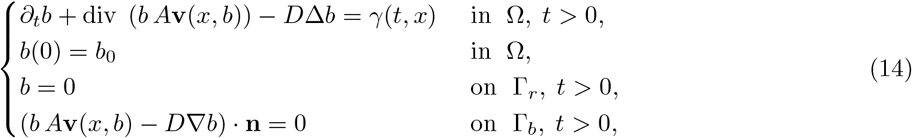

and a given *ν*(*x*_1_, *x*_2_) = (−1, 1). Since the analytical formulation of the solution is available, we impose *b*(*t, x*_1_, *x*_2_) = cos(*x*_1_) sin(*x*_2_)*e*^*t*^ for (*x*_1_, *x*_2_) ∉ Ω to account for the non-local term outside the domain. The detailed expression for *γ*(*t, x*) in (14) is given in Appendix. The numerical scheme (10) with the source term *γ* then reads as

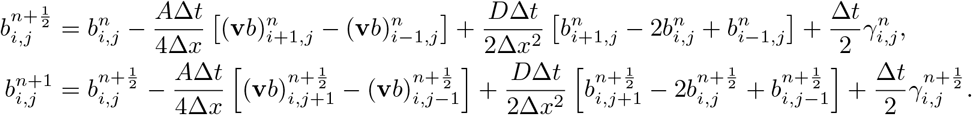

Numerical simulations of (14) were performed using the parameters specified in Table 1. The numerical order of convergence is determined as

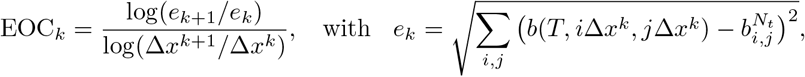

where *e*_*k*_ is the *L*_2_-error associated with the spacial step size Δ*x*^*k*^.

**Table 1:** First two columns: Parameters used in numerical simulations of problem (14), to determine the numerical order of convergence. Last four columns: Numerical *L*^2^-error and experimental order of convergence (EOC) for four refinements of the mesh.

| Name | Value | | $N$ | $\Delta x$ | $L_2$ error | EOC |
| --- | --- | --- | --- | --- | --- | --- |
| $T$ | 1 | | 32 | 0.3925 | 0.6816 | 1.0558 |
| $A$ | $\sqrt{2}/2$ | | 64 | 0.1963 | 0.3278 | 1.0186 |
| $D$ | 0.1 | | 128 | 0.0981 | 0.1618 | 1.0036 |
| $\kappa$ | 0.9 | | 256 | 0.0490 | 0.0807 | 1.0013 |
| $R$ | 1 | | 512 | 0.0245 | 0.0403 | - |
| $\lambda_{\text{CFL}}$ | 0.9 | | | | | |

The numerical scheme (10) yields the approximation for solutions to problem (14) of the first order of convergence, see Table 1 and Figure 3. We observed a non-trivial behaviour of numerical solutions of (14), with a notch visible at the centre of local extrema, see Figure 4. It was caused by the approximation of the non-linear term, which vanished when considering finer meshes.

**Fig. 3:**
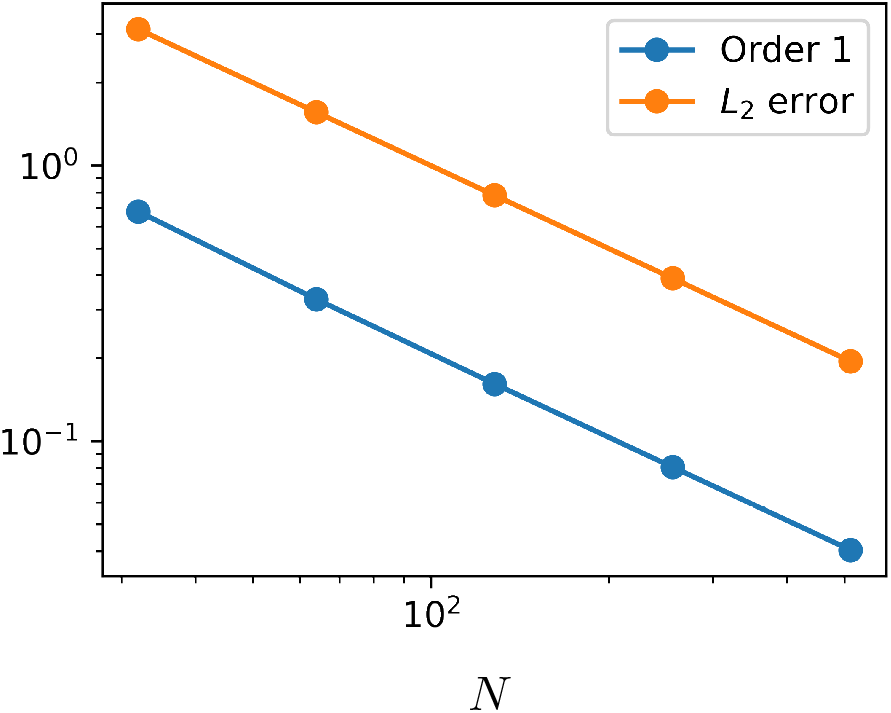
Numerical *L*^2^-error for scheme (10) for an increasing number of grid points *N*.

**Fig. 4:**
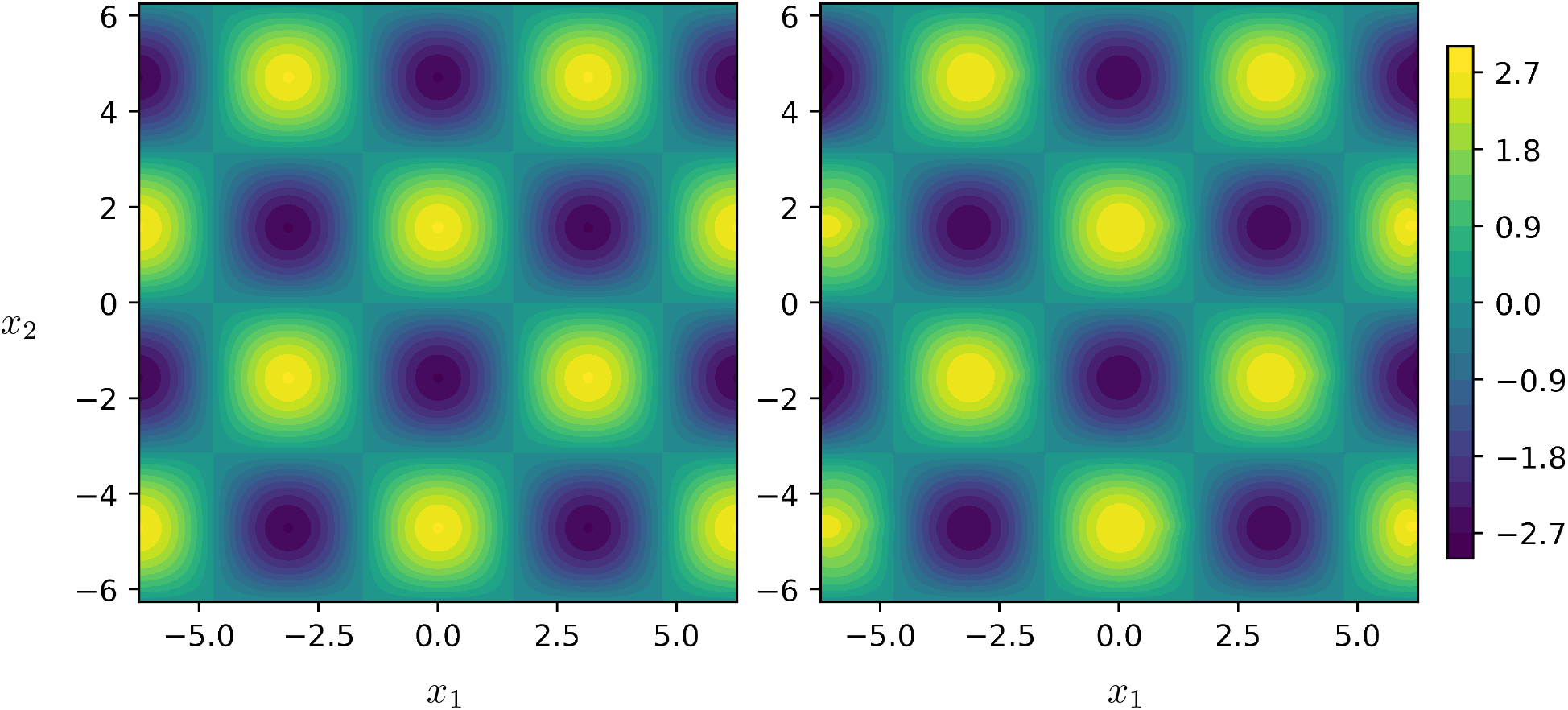
Numerical solution of (14) at *T* = 1 for Δ*x* = 0.049 and Ω = (−2*π*, 2*π*) (2*π*, 2*π*). A notch at the right of each yellow peak was observed as the result of a numerical artifact that disappears when refining the meshes.

## 4 Numerical simulations

The formation of filamentous flocks resulted from the combined effect of diffusion *D*, strength *κ* and radius *R* of interaction, but also the topology of the pore network. We performed a systematic study of how these parameters affect the size and shape of the flocks. We considered

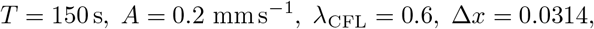

and the values for *D, κ* and *R* are shown in Table 2.

**Table 2:** Parameter values considered in numerical simulations of model (1).

| Case name | Label | $D$ | $\kappa$ | $R$ | $Ro$ |
| --- | --- | --- | --- | --- | --- |
| Reference | **R** | 0.02 | 0.5 | 1.0 | - |
| High Diffusion | **HD** | 0.1 | 0.5 | 1.0 | - |
| Low Diffusion | **LD** | 0.005 | 0.5 | 1.0 | - |
| Strong Interaction | **SI** | 0.02 | 0.9 | 1.0 | - |
| Zero Interaction | **ZI** | 0.02 | 0.0 | 1.0 | - |
| Large Radius | **LR** | 0.02 | 0.5 | 4.0 | - |
| Small Radius | **SR** | 0.02 | 0.5 | 0.1 | - |
| Tight Obstacles | **TO** | 0.02 | 0.5 | 1.0 | 3.0 |
| Medium Obstacles | **MO** | 0.02 | 0.5 | 1.0 | 6.0 |
| Far Obstacles | **FO** | 0.02 | 0.5 | 1.0 | 9.0 |

The domain Ω = [−8.0, 8.0] × [−13.0, 5.0]\Ω_0_, where 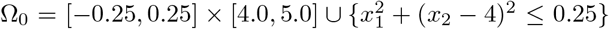 represented the tip of a root. Obstacles were represented as discs of radius 1.0 distributed on a hexagonal lattice with varying inter-particle distance *Ro*. At *t* = 0, we considered the bacterial cell density to be uniformly distributed and such that *b*_0_ = 15. The maximal computation time was *T* = 150, however, in most simulations all bacteria were attached to the root by *t* = 80, resulting into zero cell density in Ω.

We defined 10 cases with the corresponding values of *D, κ*, and *R* described in details in Table 2. We defined **R** as the reference case which dynamics can be seen on Figure 5; two cases **HD** and **LD** considered high and low values for the diffusion coefficients; two cases **SI** and **ZI** considered the impact of interaction strength on the dynamics; two cases **LR** and **SR** considered large and small interaction radius; and in the last three cases **TO, MO** and **FO** different distributions of obstacles were considered.

**Fig. 5:**
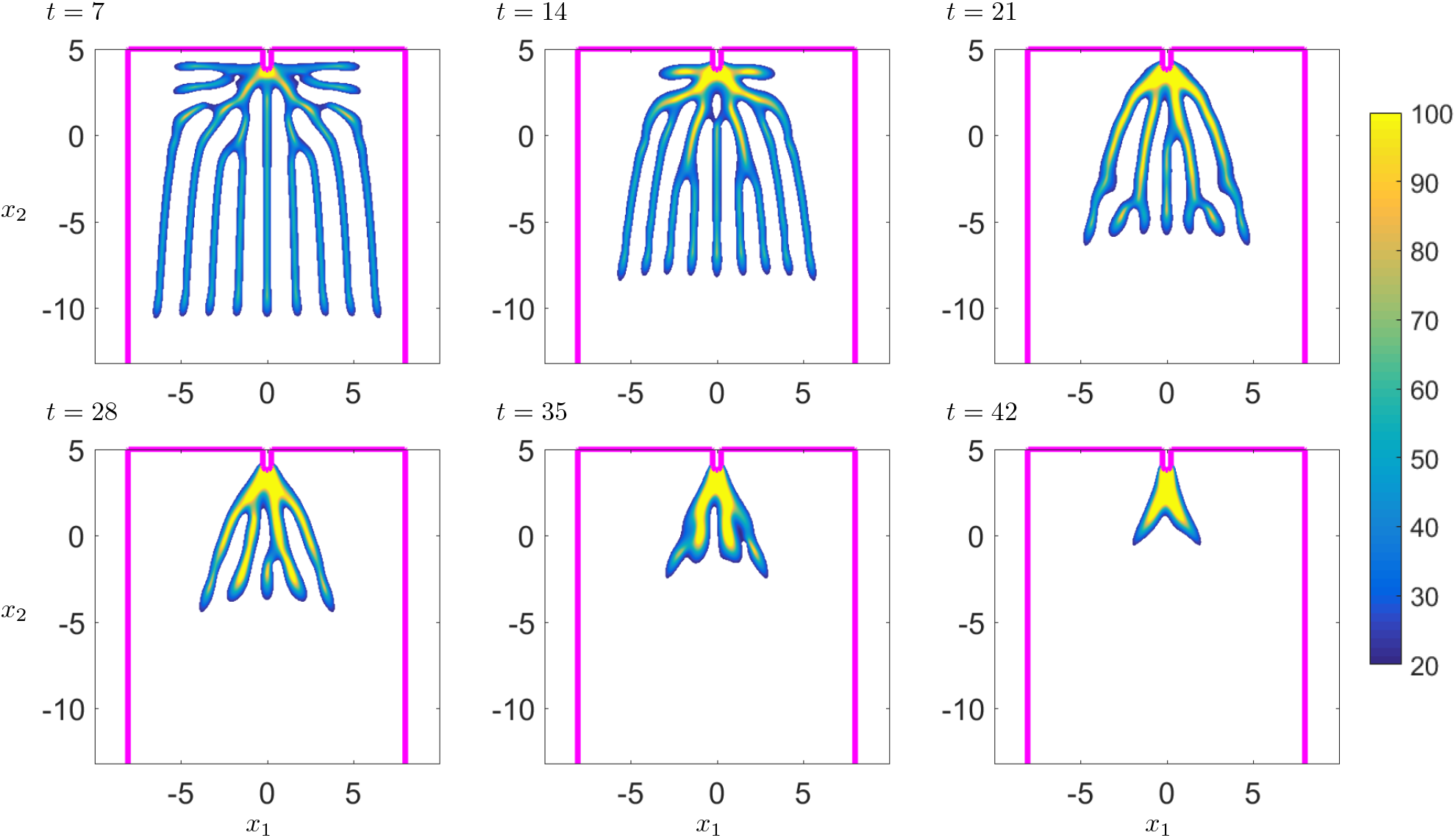
Numerical solution for model (1) taken at several time steps, with parameters for the reference case **R** defined in Table 2. The boundary *∂*Ω is traced in magenta and only bacterial cell density above 20 is plotted. Many branches formed before merging as they move towards the root.

In the case **R**, the filamentous flocks formed branches quickly with the bacteria going towards the root whilst deviating from their path as a result of peer attraction. Since no growth was considered in the model, all bacteria were attached to the surface of the root at the final time.

The diffusion coefficient *D*, the interaction strength *κ*, and the interaction radius *R* had a strong effect on the dynamics of the shape of filamentous flocks in the absence of obstacles, see Figure 6. In the cases **HD, ZI**, and **LR**, the diffusion dominated all other factors, resulting in a reduced number of branches and/or increased branch thickness. With the decrease of the diffusion coefficient, branches formed more easily, but the case **SI** showed that sufficiently large interaction strength can outweigh the attraction of the root and generate self-attracting islets of bacteria. Finally, the **SR** case showed that the interaction radius is directly related to the number of filaments and their thickness.

**Fig. 6:**
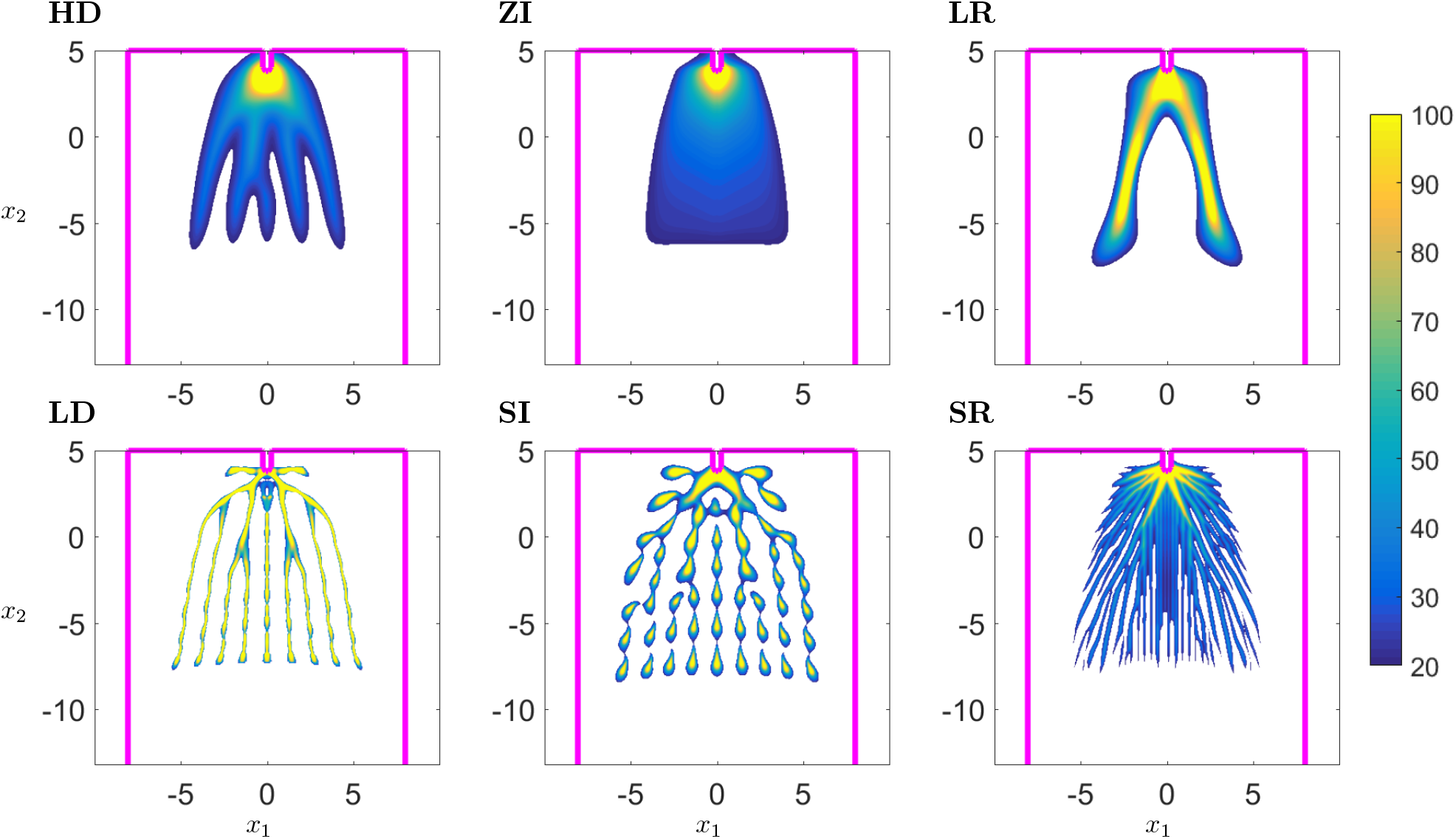
Effects of diffusion coefficient *D*, interaction strength *κ*, and interaction radius *R* on formation of filamentous flocks in the absence of obstacles. Numerical solutions of model (1) at time *t* = 14, with parameters described in Table 2. The boundary *∂*Ω is traced in magenta and only bacterial cell density above 20 is shown.

Obstacles constrained the formation of filaments through space exclusion, see Figure 7. In the case **TO**, tightly packed obstacles did not allow for variations in the formation of filaments. The presence of obstacles excluded the bacteria from some regions of the domains and led to the formation of branches. The flocking patterns were different compared to simulation results obtained in the case without obstacles **MO**. When the distance between obstacles increased, e.g. in the case of **FO**, the shapes of flocks looked more like the reference case **R**.

**Fig. 7:**
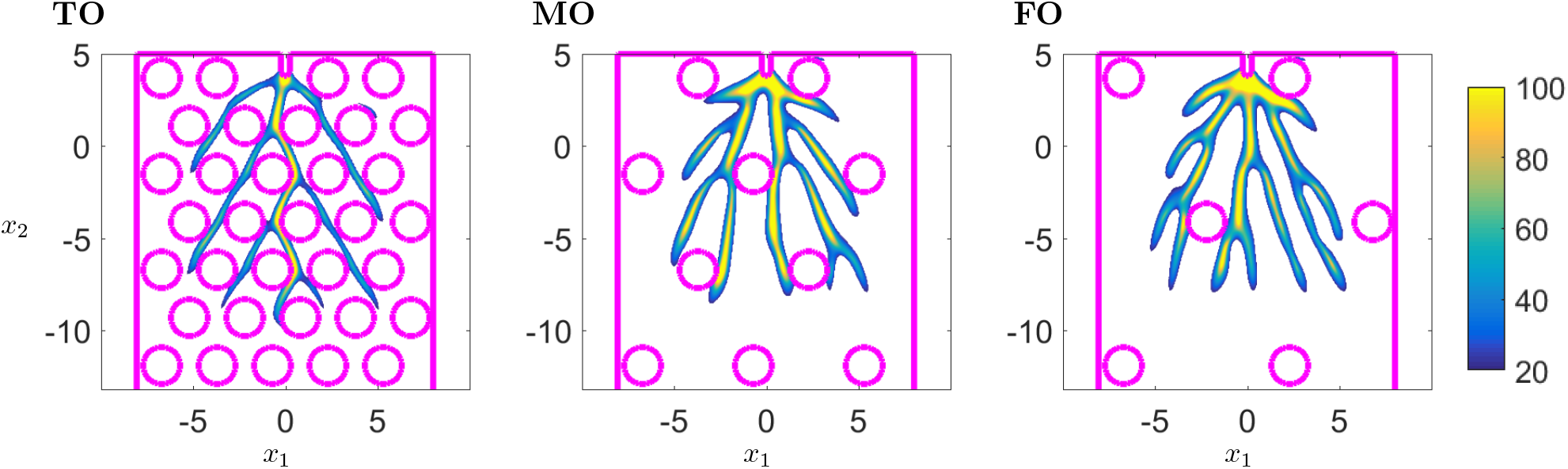
Effects of obstacle distribution on formation of filamentous flocks. Numerical solutions for model (1) at time *t* = 14 with parameters as in Table 2. Boundaries *∂*Ω are traced in magenta and only bacterial cell density above 20 is shown.

## 5 Discussion and Conclusion

We have proposed a new approach to analyse the flocking patterns of bacteria in soil. In contrast to the classical Keller-Segel equations, the chemotactic gradient here depends on the position of the plant root and on the distribution of soil particles in the domain [27, 28]. This assumption reduces the number of model parameters and allows to focus on the study of interactions between bacteria. The interactions are modeled by a non-local term with two parameters affecting peer attraction, namely the interaction strength and the interaction radius. The introduction of a non-local term unlocks new features and complex behaviours but is more challenging to implement, especially in the case of heterogeneous domains and properties. More complex forms of interactions could be introduced, e.g. nonlinear dependency on bacterial cell density [29], although experimental data are currently lacking to support such developments.

The assumption underlying the model is that the movement of bacteria is fast in comparison to the depletion of nutrients induced by the growth of bacteria. This hypothesis is reasonable because the time needed for a flock to be observed is less than 1h, which is well under the expected doubling time of the bacterium in soil [7]. In addition, the bacterial biomass introduced in the soil is orders of magnitude lower than that of the plant which release about 10 percent of its photosynthate into the soil [30]. The diffusivity of many rhizodeposits is also reduced due to their reaction and adsorption to soil particles [13]. Therefore, it is plausible that the soil chemical landscape is not significantly modified during the formation of a flock. However, the model is not adequate to describe the long term behaviour of bacterial populations when depletion of nutrients and interactions with roots occur.

In this study, we demonstrated the existence and uniqueness of solutions to the non-local model for bacteria interactions and proposed a scheme for numerical simulations of model equations, based on the finite difference and fast sweeping methods. We showed that the expected numerical order of convergence for our scheme is 1, but higher-order schemes can be implemented in the future to improve the accuracy of solutions [19].

Numerical simulations showed that the nature of interactions in bacterial populations is critical for the formation of flocks that migrate towards the plant root. The radius of interaction, which is determined by the diffusivity of quorum sensing signals, increases the diameter of the flocks. In a natural environment, flocking may reduce the energy needed for movement [31, 32]. A larger radius of interaction may therefore be beneficial when bacterial cell density is low to ensure a critical density is reached in the flock. The strength of the response to the signal from neighbouring bacterial cells also affects the dynamics of the flocks. The interaction strength controls the continuity of the bacterial flock and the ability to maintain contact with the plant root. Disconnected clusters of bacterial cells are formed when interaction strength is large while in the case of low interaction strength bacteria fail to form flocks. The diffusivity of extracellular signals, however, is affected by the environment (soil water content, viscosity) on which bacteria have little control. We can speculate, therefore, that to maintain coherent flock dynamics the sensitivity of the bacteria to a quorum sensing signal may be regulated in response to the properties of the physical environment.

## Acknowledgments

This work was funded by the European Research Council (ERC) under the European Union’s Horizon 2020 research and innovation program (Grant agreement No. 647857-SENSOILS). We also acknowledge the funding from the Spanish Ministry of Science and Innovation (MICINN) under the project MICROCROWD (PID2020-112950RR-I00). The James Hutton Institute & Scotland’s Rural College were supported by funds from the Rural and Environment Science and Analytical Services Division of the Scottish Government.

## Declarations

All authors contributed equality to the work presented in this manuscript. The code for numerical simulations are stored on the GitHub repository and will be made freely available after the manuscript is accepted for publication. There no conflicts of interest.

## Appendix

To determine *γ* in (14), we compute

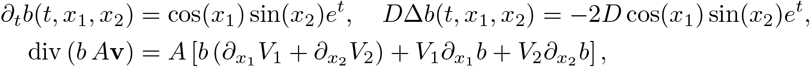

where the terms in div (*b A***v**) are given by

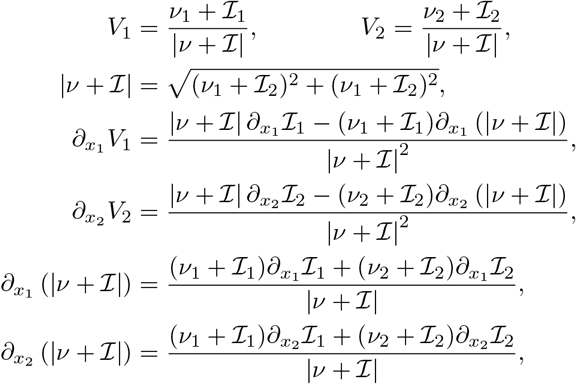

and

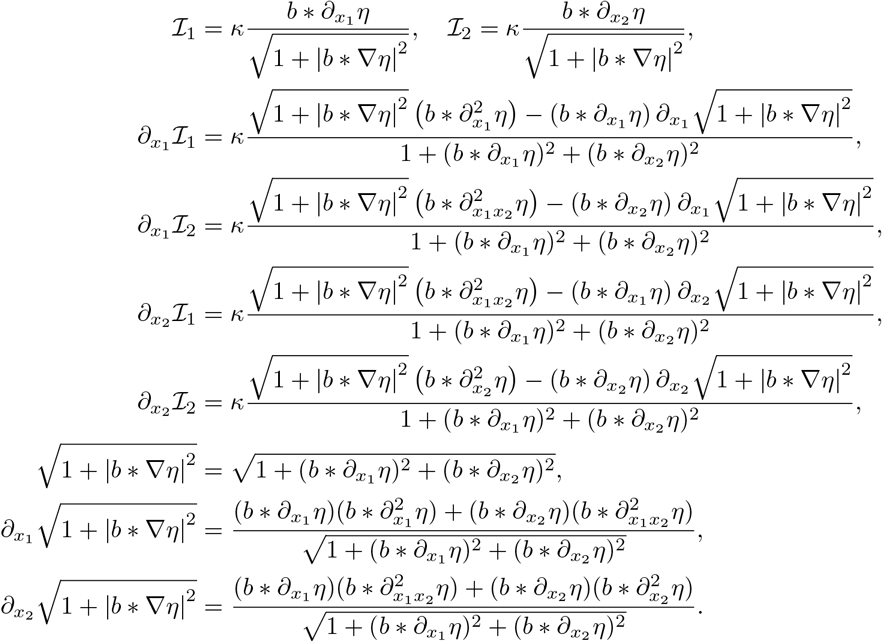

